# Rare variants association testing for a binary outcome when pooling individual level data from heterogeneous studies

**DOI:** 10.1101/2020.04.17.047530

**Authors:** Tamar Sofer, Na Guo

## Abstract

Whole genome and exome sequencing studies are used to test the association of rare genetic variants with health traits. Many existing WGS efforts now aggregate data from heterogeneous groups, e.g. combining sets of individuals of European and African ancestries. We here investigate the statistical implications on rare variant association testing with a binary trait when combining together heterogeneous studies, defined as studies with potentially different disease proportion and different frequency of variant carriers. We study and compare in simulations the type 1 error control and power of the naïve Score test, the saddlepoint approximation to the score test (SPA test), and the BinomiRare test in a range of settings, focusing on low numbers of variant carriers. We show that type 1 error control and power patterns depend on both the number of carriers of the rare allele and on disease prevalence in each of the studies. We develop recommendations for association analysis of rare genetic variants. (1) The Score test is preferred when the case proportion in the sample is 50%. (2) Do not down-sample controls to balance case-control ratio, because it reduces power. Rather, use a test that controls the type 1 error. (3) Conduct stratified analysis in parallel with combined analysis. Aggregated testing may have lower power when the variant effect size differs between strata.

## Introduction

Genetic association studies test the association of genetic variants with a trait. Genome-wide association studies (GWAS) typically test the association of each of single, common, genetic variants across the genome. This is often also done in Whole Genome Sequencing (WGS) studies, that also test rarer genetic variants. In a few examples from the WGS analysis in the Trans-Omics of Precision Medicine (TOPMed) program, investigators used a minor allele frequency threshold (MAF) of 0.001 and allowed for a minimum of 20 minor allele counts for consideration of a variant in association analyses with glycated hemoglobin (Sarnowski et al., 2019); a MAF threshold of 0.001 corresponding to at least 32 counts of the rare variant allele was applied in a study of lipids (Natarajan et al., 2018); and variants with 10 counts of the rare allele in the sample were considered in an analysis of brain imaging measures (Sarnowski et al., 2018). In other examples, investigators test rare variants associations when studying a specific gene region of interest (e.g. Amininejad et al., 2018; Tuijnenburg et al., 2018).

It is known that tests of the association of a genetic variant with a binary outcome do not control the type 1 error in some settings, and the problem is exacerbated when the genetic variant is rare (Ma et al., 2013). Specifically, when the proportion of cases in the study is low, p-values of likelihood-based tests are not well calibrated. A few tests were developed for the association of single genetic variants, that can also adjust for covariates. The Firth test (Wang, 2014), highlighted by Ma et al. (2013), uses a higher order approximation to the likelihood to compute standard errors, and is more well calibrated compared to standard tests. Rounak Dey et al. (2017) developed the saddlepoint approximation (SPA) to the p-value computation of the Score test based on a cumulant generating function rather than the standard normal distribution approximation, which is better calibrated and has improved control of type 1 error compared to the traditional Score test p-value, and is faster than the Firth test. Lee et al. (2016) developed a resampling method for calibrating single-variant tests (as well as variant-set tests), which can also account for covriates. Sofer (2017) introduced the BinomiRare test, which is robust to low case proportion and controls the type 1 error for any number of rare allele carriers. In an extensive simulation studies, Ma et al. (2013) demonstrated that the count of the rare allele determines the type 1 error and the power of statistical tests. Ma et al. (2013) and Sofer (2017) considered settings with one or multiple samples with different case proportions, however, they did not consider the scenario in which multiple samples with different frequencies of the genetic variant allele are pooled. This scenario is important, because modern large sequencing studies such as the NHLBI’s Trans-Omics in Precision Medicine (TOPMed) and the NHGRI’s CCDG aggregate individual level data from WGS studies conducted in diverse populations, where allele frequencies often differ between populations.

We set out to study rare variant association testing when pooling individual level data from various studies with potentially different population characteristics: allele frequency and case proportion. To limit the high number of possible combinations of studies’ characteristics, we focus on two studies of the same sample size and vary the disease prevalence in each of the studies as well as the count of the rare variant allele. We focus on tests that can account for covariates, and that do not use resampling, to limit computation time.

## Methods

### Logistic association model for two studies

Suppose that individual level data from two studies with *n*_1_ and *n*_2_ individuals respectively are combined. For study *j* ∈ {1,2} Let the binary outcome *D*_*ij*_ ∈ {0,1}, *i*_*j*_ = 1, …, *n*_*j*_ follow a logistic model with

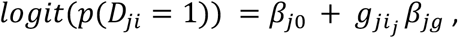

here assuming no confounders or covariates are adjusted for. When the data are pooled across studies, the model can be written instead as

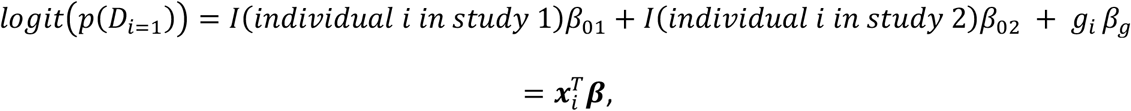

where we now add study-specific intercepts in the joint model. Note that this formulation is statistically equivalent to a formulation with the same intercept for all individuals, and a covariate for one of the studies. To simplify exposition, let ***x***_*i*_ = (*x*_*i*1_, *x*_*i*2_, *g*_*i*_)^*T*^, ***β*** = (*β*_01_, *β*_02_, *β*_*g*_)^*T*^.

### Tests for association of a variant with the outcome

Both the Score and the BinomiRare tests (and the SPA, which is a score test with better calibrated p-value) use estimates of within-sample disease probabilities under the null hypothesis of no association between the outcome and the genetic variant, i.e. under *H*_0_: *β*_*g*_ = 0. Clearly, under the null, 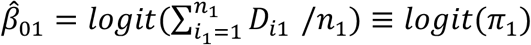, and 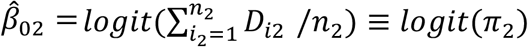, where *expit*(*x*) = *exp*(*x*)/[1 + *exp*(*x*)] is the inverse function of the logit function. The derivative of the *expit*(·) function is 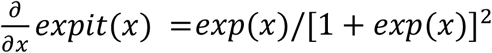. For *n* = *n*_1_ + *n*_2_, the score for *β*_*g*_ is derived as:

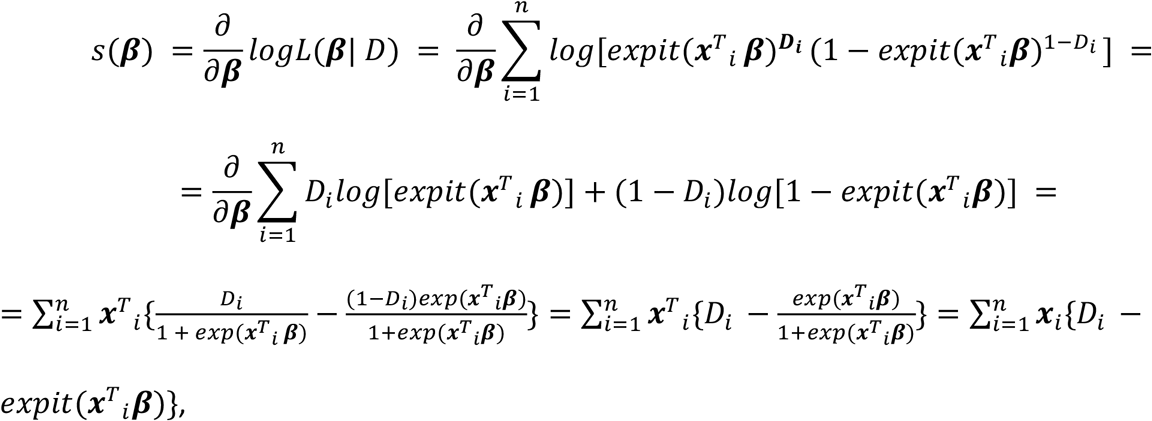

where in the score test for *β*_*g*_, *β*_01_ and *β*_02_ are estimated under the null, and in this setting, the score for *β*_*g*_ simplifies to:

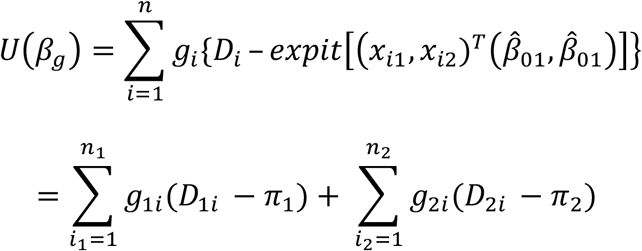

If a genetic variant is rare, then most carriers of the variant are heterozygotes, i.e. most people have *g*_*i*_ = 0, a few people have *g*_*i*_ = 1, and almost no one has *g*_*i*_ = 2, meaning that we can assume a dominant mode of variant association. We then further simplify this expression by introducing additional notation. For study *j*, let *c*_*j*_^0^ and *c*_*j*_^1^ be the number of carriers of a rare variant allele among people with the outcome *D*_*ij*_ = 0 and among those with the outcome *D*_*ij*_ = 1, respectively. Then the score is now:

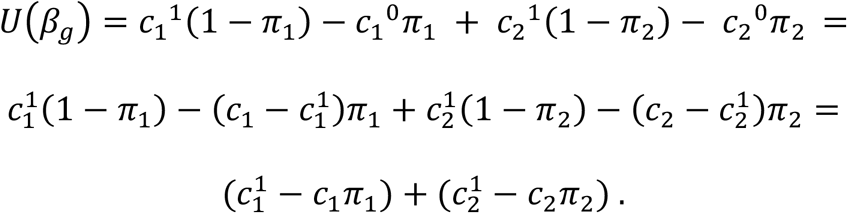

The score for *β*_*g*_ is the sum of scores in each of the two studies, and in each study, the score is a difference between the observed and the expected number of diseased carriers, under the observed disease proportion in the study.

In the standard Score test, the variance of the score for *β*_*g*_ is estimated by deriving the information matrix, and then extracting the appropriate entry from its inverse. For logistic regression, the information matrix is given by:

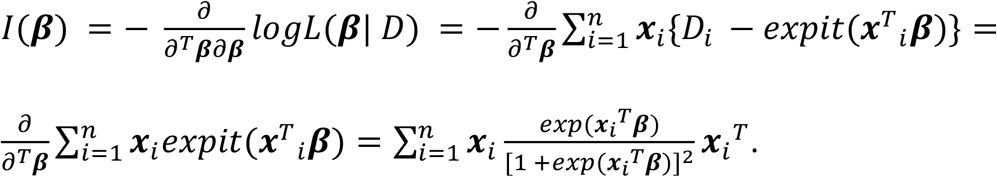

This can be written in a matrix form. Define the following matrices:

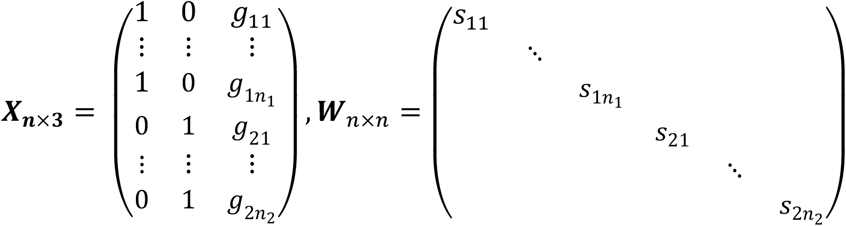

where ***X*** is the design matrix of the regression of the disease on the variant, accounting for two studies using study-specific intercepts, and ***W*** is the diagonal matrix with diagonals, for person *ji* from study *j, j* ∈ {1,2}, *i* = 1, …, *n*_*j*_ having 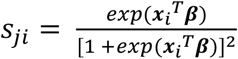. Then

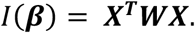

Noting that for the Score test, the information matrix will be evaluated under the null; therefore, the only covariates are the study-specific intercepts, we have that for all individuals in study one 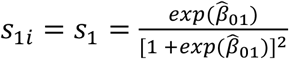, and in study two 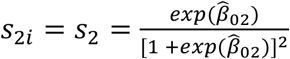 Then:

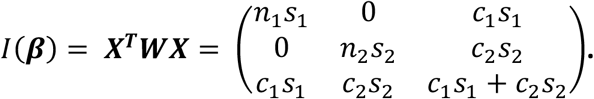

Using formula for matrix inverse, one can compute the entry of *I*(***β***)^−1^ corresponding to *β*_*g*_, as

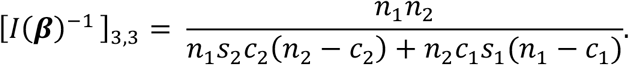

Its inverse is the variance of the score:

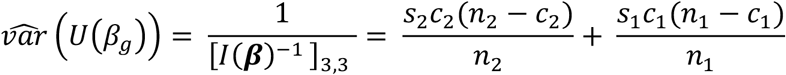

which is the sum of the scores for *β*_*g*_ in each of the two studies. The estimator of the variance of the score depends, through *s*_1_, *s*_2_, on the observed outcome proportion in the sample, on the observed variant allele count in the sample, and on the sample size. When the observed variant count is very low compared to the number of individuals in the study, e.g. when *c*_1_ is fixed and *n* → ∞, we have that *c*_*j*_(*n*_*j*_ − *c*_*j*_)/*n*_*j*_ ≈ *c*_*j*_ for *j* ∈ {1,2}, so that both the score and its variance do not depend on the sample size, but rather only on the variant count and the disease proportion in the study. Note that this asymptotic setting is standard for genome sequencing data because as the sample size grows more low-count variants are observed.

The SPA test (R. Dey et al., 2017) instead of using the above score variance estimates, computes a p-value based on a obtaining a better approximate distribution of the test statistic using a cumulant generating function, and uses the saddlepoint approximation to solve the resulting optimization problem.

The BinomiRare test (Sofer, 2017) only relies on the observed outcome frequency (more generally, the outcome probabilities) in the carriers. It takes the vector of estimated outcome probabilities for the carriers of the rare variants, and uses the Poisson-Binomial distribution (Hong, 2013) to compute a p-value for testing the null hypothesis of no association between the variant and the outcome. Therefore, in our simplified settings that do not use covariates, the BinomiRare tests depends on the numbers of carriers *c*_1_, *c*_2_, diseased carriers 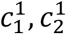, and the proportions of diseased individuals in the studies. Because the BinomiRare test does not use a normal approximation to the Poisson-Binomial distribution, it has a discrete probability mass function. Two types of p-values can be computed: the standard p-value, and the mid-p-value. Let 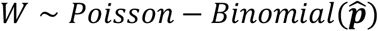, with 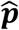 being the vector of estimated disease probabilities for the *c*_1_ + *c*_2_ carriers of the rare-variant of interest in the two studies. The p-value and mid-p-value are defined as:

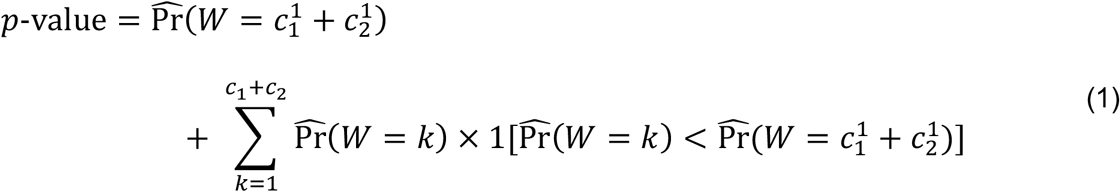

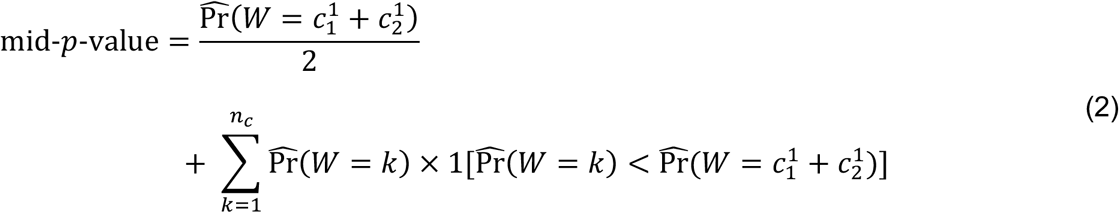

Clearly, the mid-p-value (2) is less conservative, and therefore will always be smaller than the p-value (1). However, when the number of carriers is small, it may be too liberal.

### Simulation studies

In our simulation studies described henceforth, we generated datasets in the simple settings described above, and used the computationally efficient implementation of the Score test *U*(*β*_*g*_) and 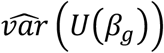. For the SPA test, we used the naïve Score test p-value. When it was smaller than 0.05, we re-computed a p-value using the SPAtest R package (Dey & Lee, 2018).

### Simulation studies: type 1 error control when combining two studies

We considered four settings of rare variant distributions across studies: (*c*_1_, *c*_2_) = {(10,10), (100,100), (10,100), (100,10)}. We varied the disease proportions in each of the two studies, so that the disease proportion in study 1 was *expit*(*β*_10_) ∈ {0.01, 0.05, 0.2, 0.5}, and the disease proportion in study 2 was taking the same values across the simulation studies, so that it was always lower or equal to the proportion in study 1. In each of the settings defined by number of carriers and disease proportions we performed 10^8^ simulations with *n*_1_ = *n*_2_ = 10,000, and evaluated type 1 error when using the Score, SPA, and BinomiRare tests with a p-value threshold of 10^−4^. For the BinomiRare test, we used both the p-value (equation (1); pval) and the mid-p-value (equation (2), midp). For comparison, we also studied the following settings:

1. A simulation study with *n*_1_ = *n*_2_ = 5,000, holding all other parameters the same.
2. A simulation study in which we plugged-in the true, known, per-study disease prevalence *expit*(*β*_10_), *expit*(*β*_20_) rather than estimated them when computing the Score statistics, with provided disease probabilities for computing BinomiRare mid-p-values.
3. Simulation studies in which we generated datasets by down-sampling controls from each study, with a ratio of up to 3 controls per case and with 1 control per case.

Finally, we also performed additional simulations for some of the “baseline” scenarios (n_1_ =n_2_=10,000, no down-sampling) in which we expended the number of simulation repetitions to 10^10^, to estimate type 1 error at a range of p-value threshold for testing, with the lowest value being 5×10^−8^.

### Simulation studies: identifying minimum number of carriers for SPA test

We performed a simulation study with the goal of formulating a recommendation for the minimum number of carriers required for appropriate type 1 error control by the SPA test when combining individual level data from heterogeneous studies. To limit the potential number of simulations, we focused on three possible settings of disease proportion in the two studies: [*expit*(*β*_10_), *expit*(*β*_20_)] ∈ {(0.01, 0.01), (0.01, 0.5), (0.5, 0.5)}, and the number of carriers in the two studies was varied so that all possible combinations of *c*_1_, *c*_2_ ∈ {10, 15, 20, 25, 30} were evaluated. We also studied the same settings with the BinomiRare test, with both the pval and midp options. For the SPA test only, we considered additional settings in which *c*_1_ = *c*_2_ ∈ {35, 40, 45, 50, 55}.

### Simulation studies: power comparisons

To study the implications of testing genetic variants in an aggregated dataset of two heterogeneous studies, and to compare power between tests, we used similar settings to those used for type 1 error assessment, with the same follow-up comparisons. To generate datasets for these simulations, we allowed for different probability of disease among carriers of the rare variant, so that 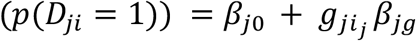. First, focusing on tests comparison, sample size, and down-sampling of controls, we used the same effect size *β*_1*g*_ = *β*_2*g*_ = *β*_*g*_ in the two studies combined together. Effect sizes varied *β*_*g*_ ∈ {log(2), log(3), log(4), log (5)}. Next, we studied the implication of having different effect size in the two studies, while setting *β*_1*g*_ ∈ {log (3), log(4)} and *β*_2*g*_ ∈ {log(1), log(1.5), log(2)}, i.e. settings where the variant is causal only in study 1, and when it has a very small effect in study 2 compared to that in study 1. We used the same p-value threshold of 10^−4^ as before.

### Computing approximate power for BinomiRare test

On the dedicated GitHub repository https://github.com/tamartsi/Binary_combine we provide a function to compute approximate power for the BinomiRare test. The function takes a vector of estimated outcome probabilities in the sample under the null hypothesis of no association between genotype and outcome, an odds ratio parameter, p-value threshold for declaring significance, number of carriers ‘n_carrier’, and number of simulation iterations. Then, in each simulation iteration it uniformly samples n_carrier outcome probabilities (without replacement). For each sampled carrier, given its outcome probability under the null *p*_0_, the function computes the outcome probability under the alternative hypothesis *p*_*A*_ = *expit*[*logit*(*p*_0_) + log(*OR*)], and uses the binomial distribution to simulate outcome status using *p*_*A*_. Then, it uses the BinomiRare test to compute a p-value for the null hypothesis of 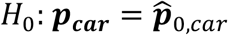, where ***p***_***car***_ is the true vector of outcome probabilities among the carriers, and 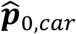, is the vector of estimated outcome probabilities under the null. Finally, the power is the proportion of p-values computed in the simulations which were lower than the p-value threshold.

## Results

### Type 1 error when combining two studies

Figure 1 provides type 1 error comparisons when combining two studies with each *n*_1_ = *n*_2_ = 10,000, with varying disease proportions in the two studies, and four scenarios of number of carriers across the studies. We compare the naïve Score test, the SPA, and the BinomiRare test with the pval (usual p-value, equation (1)) and midp (mid-p-value, equation (2)) options. The figure provides the observed test size divided by the desired type 1 error. Ideally, this number should be 1. Higher numbers indicated inflation (larger number of false detection than expected), and lower numbers indicate deflation, or conservativeness.

**Figure 1:**
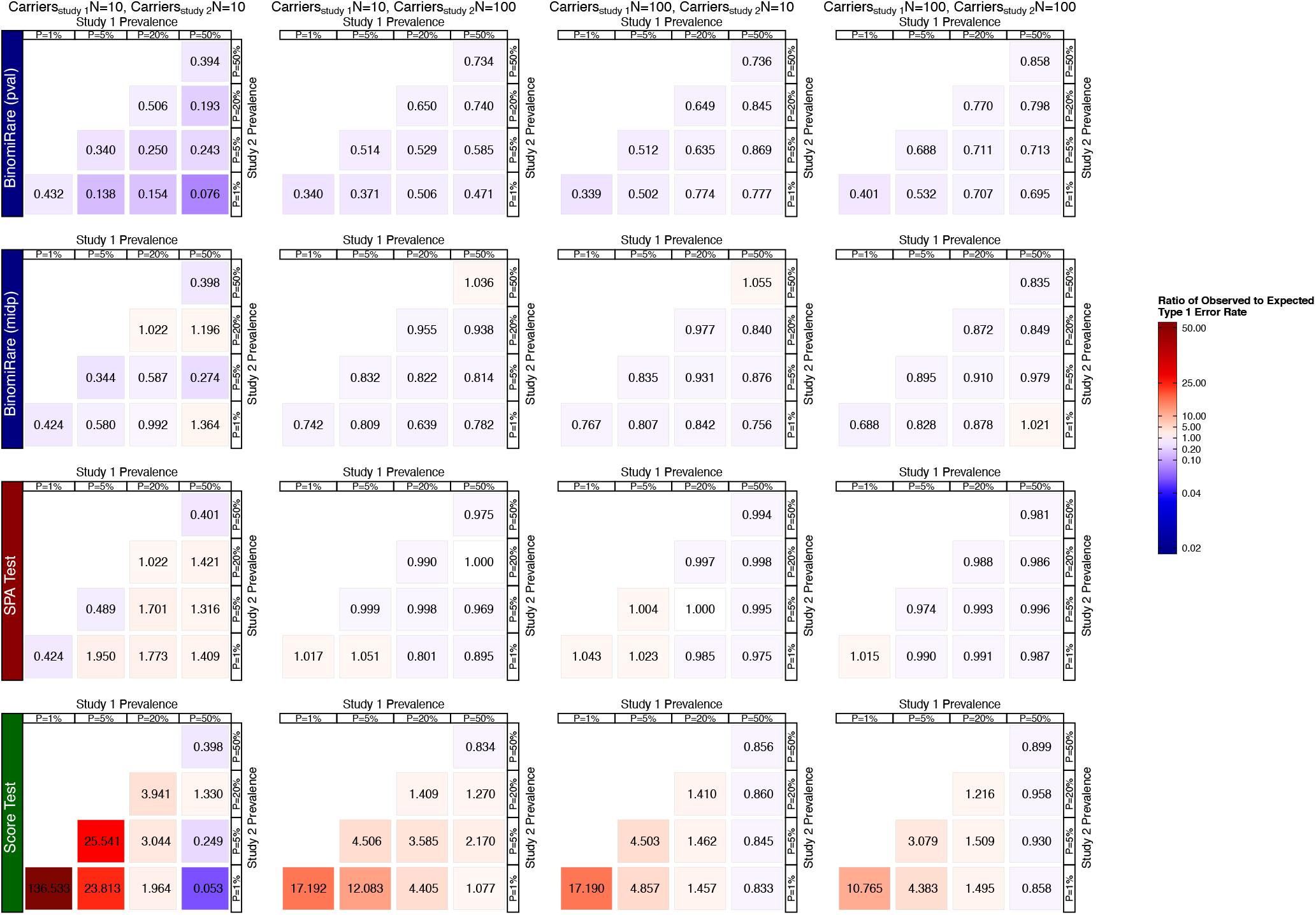
Ratios between observed and expected type 1 error rate in simulation studies when testing a binary outcome for association with a rare genetic variant. We compared the naïve score (Score) test, the SPA test, and BinomiRare with the usual p-values (pval) and the mid-p-value (midp), for settings defined by the number of carriers and outcome prevalence in each study. Both studies had n=10,000 individuals. For each setting we performed 10^8^ simulations, and the p-value threshold used for determining significance was 10^−4^. Values of 1 correspond to perfect calibration, and values larger (smaller) than 1 correspond to inflation (deflation), or higher (lower) number of detected false associations.

#### Score test

As is already known, we see that the naïve Score test becomes more inflated as the disease prevalence in the total sample becomes low. Overall the Score performance become better with increased number of carriers in the combined sample. However, for a fixed number of carriers, there is a difference in performance depending on which of the two studies have more carriers: when considering the two non-symmetric scenarios, i.e. the scenarios in which *c*_1_ = 10; *c*_2_ = 100, and the other way around, we see that the Score test performance depend on the number of carriers in each study. Specifically, in comparison with the settings of *c*_1_ = *c*_2_ = 10, if an analyst required at least 100 carriers in the study with higher disease prevalence but allowed the number of carriers in the other study to stay 10, the inflation was reduced compared to the settings in which 100 carriers were required in the study with lower case proportion. If the analyst further required both studies to have at least 100 carriers, the inflation did not improve much. This suggest that when combining multiple studies, it may be useful to require a minimum number of carriers in the study with the higher disease proportion in order to stabilize the Score test results.

#### SPA test

Type 1 error control was mostly appropriate when the total number of carriers in the combined two-study sample was 110 or 200, with a few settings with low degree of inflation (see Figure 1). When there were 20 carriers in the combined sample, type 1 error was usually not controlled, other than in the settings in which the disease proportion was equal in the study 1 and study 2. In this case, SPA was often conservative.

#### BinomiRare

Type 1 error was always controlled when the usual p-value (pval) was used, and usually controlled with the midp option. In a few settings, the BinomiRare with the midp option had low degree of inflation. Due to the discreteness of the Poisson-Binomial distribution (which is not approximated to a normal distribution by this test), the size of the test when using the pval option is often conservative.

#### Low p-value thresholds

in Supplementary Tables 1-4, we also report the ratio between observed to expected type 1 error in various settings when the p-value threshold used is as low as 5×10^−8^. SPA test often results in the desired ratio of 1 (no inflation/deflation), with the exception of low number of carriers (20 in the combined dataset), and some low grade inflation/deflation (0.8—1.3) when there were 10 carriers in study 1 and 100 in study 2, while the disease proportion was higher in study 1. BinomiRare (pval) was often very conservative (observed to expect type 1 error ratio < 1), and BinomiRare (midp) was also usually conservative, with a handful of settings with low-grade inflation. The inflation of the Score test usually increased as the p-value threshold become lower.

#### Other settings

Comparisons of some of the above simulations to settings where *n*_1_ = *n*_2_ = 5,000 show that the results are mostly the same, confirming that the properties of the tests mostly depend on the number of carriers (Figure 1 in the Supplementary Information). To address the question of whether and how the results are strongly affected by estimation of disease probabilities, which do depend on sample size, we also compared type 1 error between the main simulation study and a simulation study in which disease probabilities are taken as known for both the Score and the BinomiRare test (Figure 1 in the Supplementary Information). Type 1 error control often improved for the Score test, but in some cases became worse for the BinomiRare, when using the midp option. To note, BinomiRare was always more conservative than the Score test in this simulation. When down-sampling controls to reduce case-control ratios, Figure 2 demonstrates that the type 1 error is always controlled by the Score test if the case-control ratio is 1:1 (as expected), but not when the case-control ratio is 1:3.

**Figure 2:**
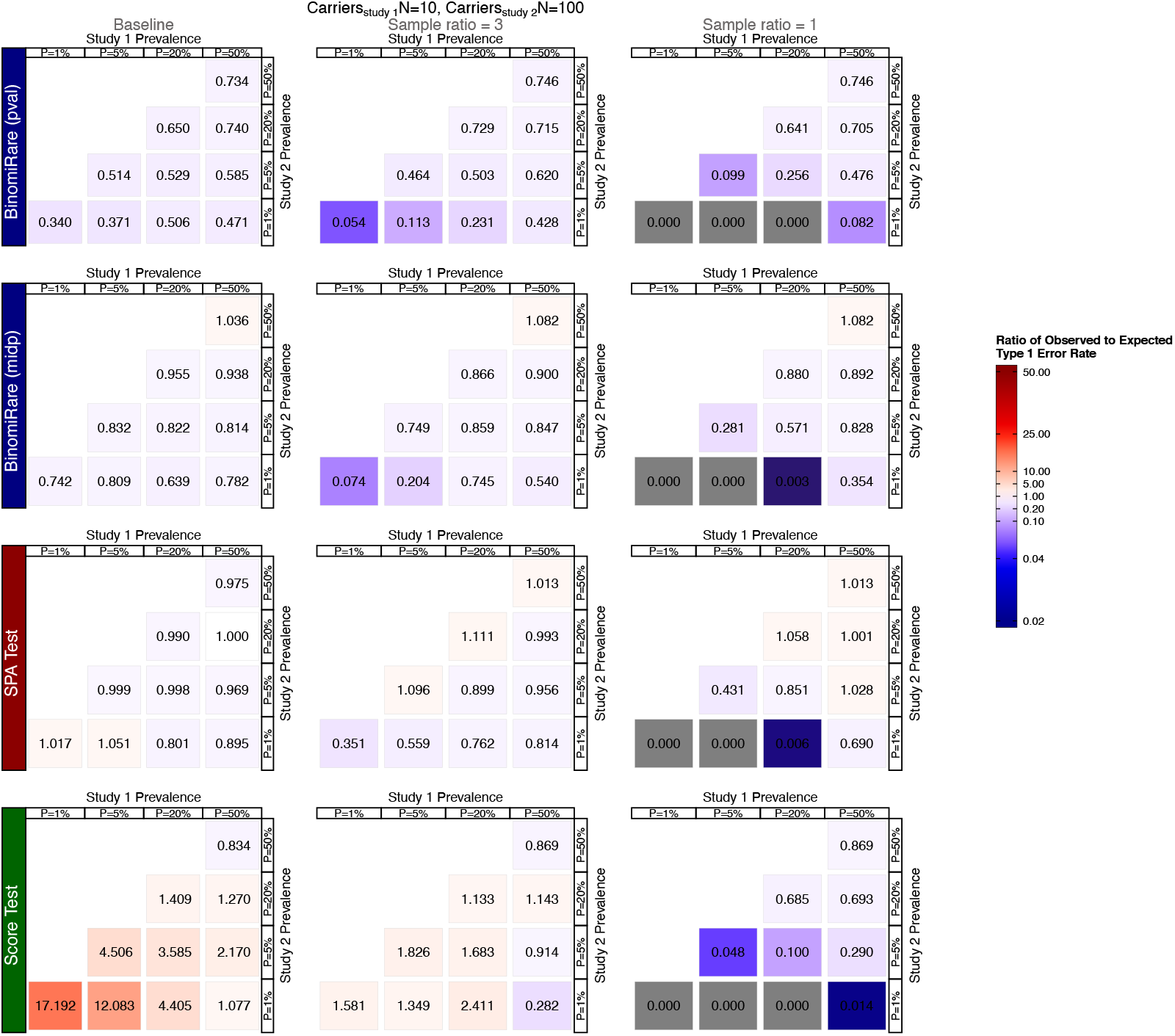
Ratios between observed and expected type 1 error rate in simulation studies when testing a binary outcome for association with a rare genetic variant. We compared the naïve score (Score) test, the SPA test, and BinomiRare with the usual p-values (pval) and the mid-p-value (midp). In all settings data from two studies were pooled together, with 10 carriers of the rare variant in study 1, and 100 carriers of the rare variant in study 2. Both studies had n=10,000 individuals. The settings investigated here are defined by the outcome prevalence in each study. The left column (“Baseline”) correspond to analysis of the complete data. The middle (“Sample ratio = 3”) and right (“Sample ratio = 1”) columns provide results for analyses that reduced sample sizes by down-sampling controls to generate samples with the specified control:case ratio. For each setting we performed 10^8^ simulations, and the p-value threshold used for determining significance was 10^−4^. Values of 1 correspond to perfect calibration, and values larger (smaller) than 1 correspond to inflation (deflation), or higher (lower) number of detected false associations.

Supplementary Figures 2 and 3 provides all settings under down-sampling of controls with ratio 1:3 and 1:1, respectively. All tests become very conservative when the total number of carriers in each of the studies is 10 prior to down-sampling controls because often no carriers are left in the analytic sample after down-sampling of controls. Further, when down-sampling controls the SPA test often becomes inflated at times, especially in the 1:1 down-sampling scenario, likely because the number of carriers remaining in the data after down-sampling of controls is very low.

### Simulations to study the minimum number of carriers for SPA

In the simulations designed to study the minimum number of carriers for SPA, which had up to 60 carriers in the combined sample, type 1 error was not perfectly controlled even when there were 60 carriers and the case proportion was 50% in both studies (Supplementary Figure 4). Because we did not see any pattern related to type 1 error control with respect to the distribution of variant carriers across studies, we also considered scenarios with an equal number of carriers in each study, with up to 55 carriers. Figure 3 provides the type 1 error for the SPA test when the number of carriers was equal in the two combined studies, and ranged from 10 to 55, by increments of 5. When the number of carriers was 45 in each study or 90 in the combined sample, the type 1 error is controlled. Then, in the setting with 55 carriers in and case prevalence of 0.01 in each study, the type 1 error was 1.14×10^−4^, which is larger than expected in the 95% confidence intervals accounting for p-value threshold of 1×10^−4^ and 1×10^8^ simulations.

**Figure 3:**
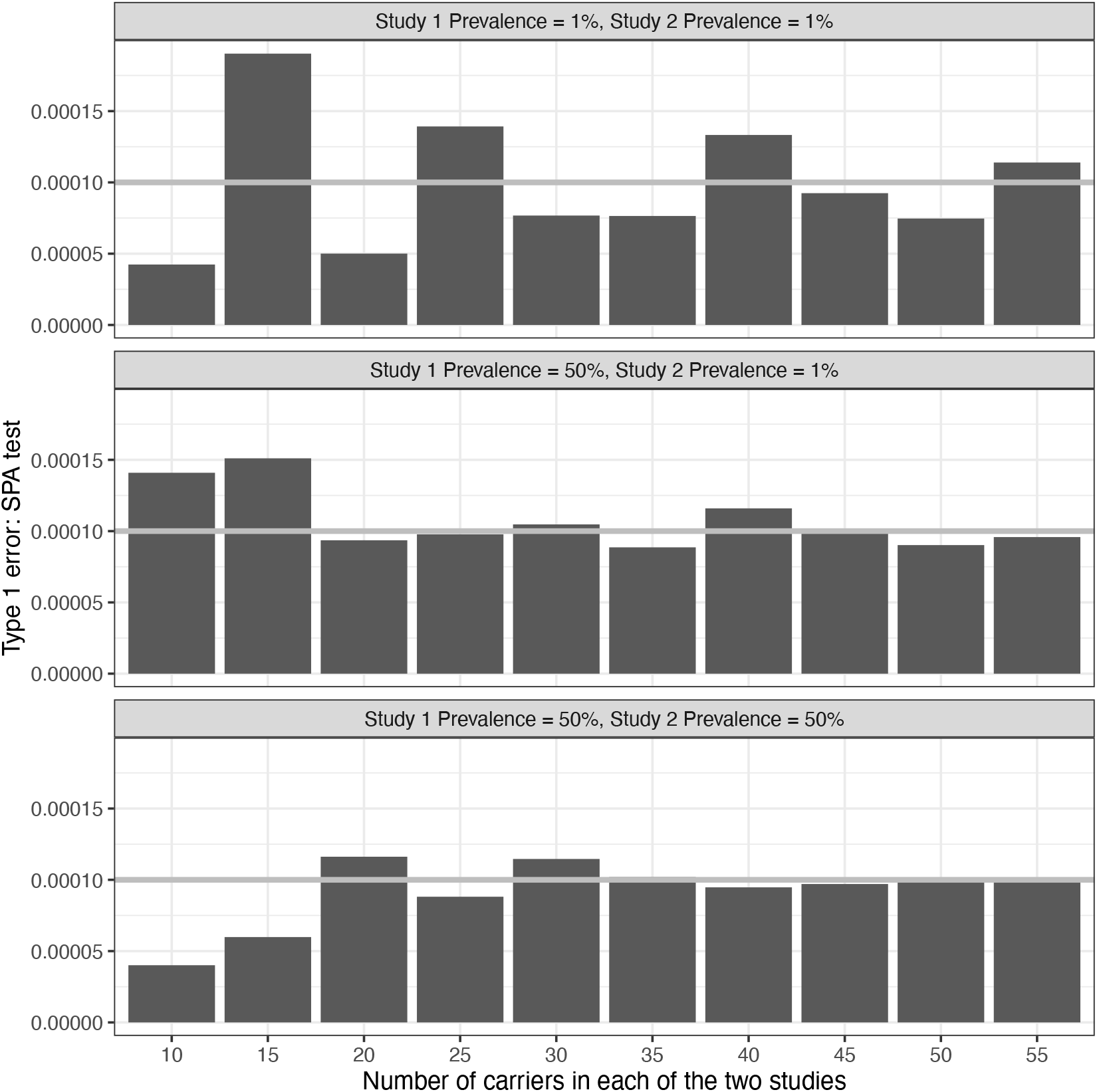
Type 1 error of the SPA test in simulations combining two studies with n=10,000 and equal number of carriers in each study. Simulation settings are defined by the prevalence of the outcome and the number of carriers in each of the studies. For each setting we performed 10^8^ simulations, and the p-value threshold used for determining significance was 10^−4^. Values of 1 correspond to perfect calibration, and values larger (smaller) than 1 correspond to inflation (deflation), or higher (lower) number of detected false associations.

### Power when combining two studies: equal variant effect size in the two studies

Figure 4 compares power between the various tests when the case prevalence was 20% in study 1, 5% in study 2, for a few carriers setting, and comparing the baseline simulations (no down-sampling of controls), and down-sampling of controls with case-control ratio of 1:3 and 1:1. The figure provides the estimated power even when tests did not control the type 1 error in the corresponding simulation studies (while highlighting this non-control). For 110 carriers in the combined sample of 20,000 people, the power is higher when there are 100 carriers in the study with 20% cases, compared with 100 carriers in the study with 5% cases. This is true in other simulations as well: for a fixed number of carriers in the total sample, power is higher when more carriers are in the study with higher case proportion. Power is reduced when controls are down-sampled, especially when the effect size is small. Among the two settings of 110 carriers in the combined sample, down-sampling of controls leads to more substantial reduction of power when the number of carriers is 100 in the study with lower case prevalence. This is likely because the down-sampling is more aggressive (lower total sample size), resulting in a substantially reduced number of carriers after down-sampling.

**Figure 4:**
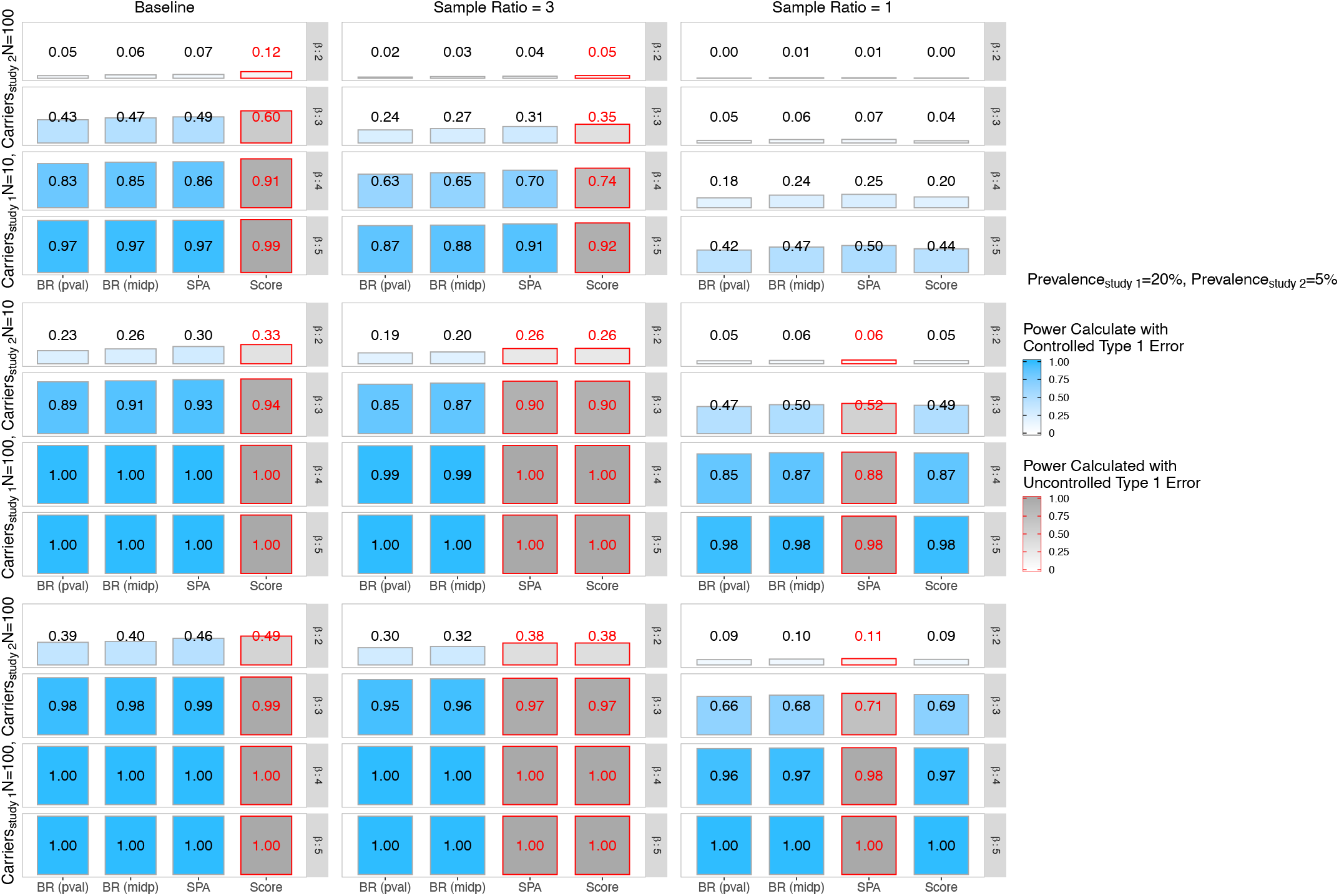
Power estimated in simulation studies when testing a binary outcome for association with a rare genetic variant. We compared the naïve score (Score) test, the SPA test, and BinomiRare with the usual p-values (pval) and the mid-p-value (midp). The simulation settings are defined by the number of carriers in each of the studies, and the variant effect size *β*. The outcome prevalence was fixed at 0.2 in study 1 and 0.05 in study 2. The left column (“Baseline”) correspond to analysis of the complete data. The middle (“Sample ratio = 3”) and right (“Sample ratio = 1”) columns provide results for analyses that studies sample sizes by down-sampling controls to generate samples with the specified control:case ratio. For each setting we performed 10^4^ simulations, and the p-value threshold used for determining significance was 10^−4^. We color coded the settings according to type 1 error control in the simulations corresponding to the same prevalence, carrier, and down-sampling settings, but with no variant association.

We also compared power in the main simulations to the settings when there were only 5,000 individuals in each study (but the same number of carriers), and when the case prevalence was known, providing true outcome probabilities as plug-ins for BinomiRare and Score tests (Supplementary Figure 5). When n=5,000 in each study, the power was about 90-100% of the power in the corresponding setting when there were n=10,000 individuals in each study, suggesting that more precisely estimated outcome probabilities could increase power. However, using the two outcome probabilities did not generally increase the power of BinomiRare, perhaps because in the simple investigated settings the parameter estimates are already quite precise.

Finally, we note that the BinomiRare with the mid-p-value is more powerful than the BinomiRare test with the usual p-value, as is known by definition, with BinomiRare-midp having up to 111% the power of the BinomiRare-pval option. The SPA test was up to 116% more powerful than the BinomiRare-midp test (focusing these comparisons on settings where both tests controlled the type 1 error).

### Power when combining two studies: one study has no or low variant effect

We performed simulations studies in which a variant was associated with the disease in study 1, and was either not associated with the disease in study 2, or had weaker association. The pattern of results was the similar across the different tests, so we provide results from BinomiRare pval, because it always controls the type 1 error and therefore the power is accurate. Figure 5 provides estimated power when there are 100 of carriers in study 1, and the disease prevalence in study 1 is 0.05, and the effect size β_1g_ = log(4), across settings of disease prevalence, odds ratios exp (β_2g_), and number of carriers in study 2. The power is compared to that when testing the variant association using study 1 only (grey line). When the variant is not associated, or has low association with the disease in study 2 (*β*_2*g*_ ∈ {log(1), log(1.5)}), aggregating individual level data from the two studies results in decreased power compared to testing in study 1 alone. The power is lower when the disease prevalence is higher in study 2, and when the number of carriers is 100, compared to 10. When *β*_2*g*_ = log (2), including study 2 in analysis increases power when the disease prevalence is high and the number of carriers is high in study 2. When disease prevalence is low (0.01), the power is the same whether study 2 is included or not. Supplementary Figure 6 provides similar results in the settings where the disease prevalence in study 1 is 0.5, and the effect size β_1g_ = log(3). In this case, there is lower loss of power from including study 2.

**Figure 5:**
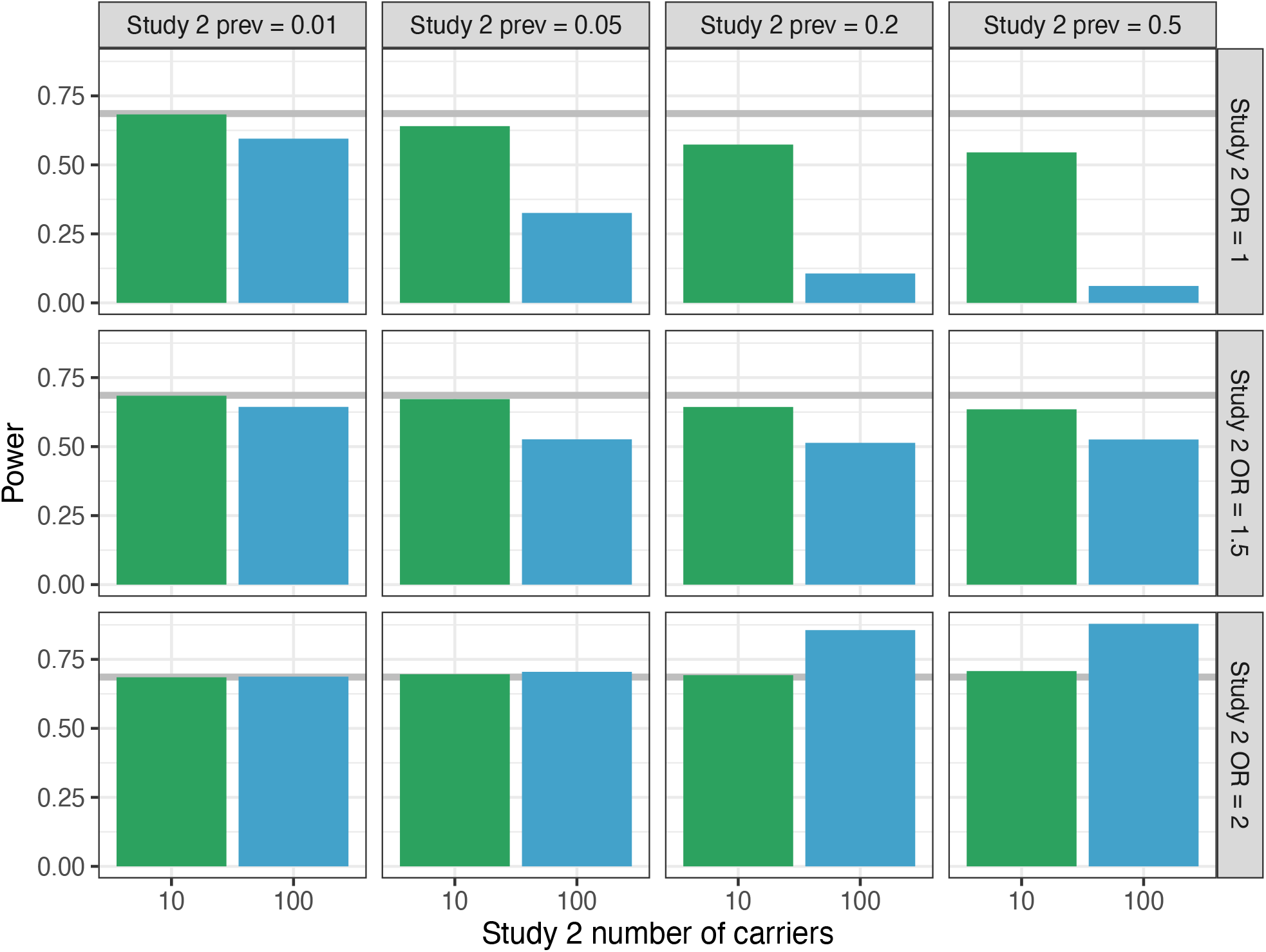
Power estimated in simulation studies when testing a binary outcome for an association with a rare genetic variant using the BinomiRare test (using the usual p-values, pval). In study 1, the outcome prevalence was always 0.05, there were 100 carriers, and the variant effect size was *β*_1*g*_ = log (3). Each block in the figure provides estimated testing power for a given outcome prevalence in study 2 (columns), odds ratio (OR) exp (*β*_2*g*_) in study 2 (rows), and for 10 and 100 carriers in study 2. The grey line corresponds to the estimated power for testing the variant association in study 1 alone. For each setting we performed 10^4^ simulations, and the p-value threshold used for determining significance was 10^−4^.

### Recommendations

We here provide analytic recommendations for investigators studying rare variants by pooling heterogeneous studies such as TOPMed.

1. If the case proportion is 0.5, use the Score test. While it sometimes controls the type 1 error also in lower case proportions, this depends on multiple factors, including the p-value thresholds (see Supplementary Tables 2 and 3), so as a rule of thumb we do not advise using it for rare variants if the case proportion is lower than 0.5.
2. When the case proportion is low, do not down-sample controls, because you will lose statistical power. Instead, use a test that controls the type 1 error such as the BinomiRare (controls the type 1 error for any carrier count, but tends to be less powerful) or SPA test (sometimes does not control the type 1 error with low carrier count, but is otherwise more powerful than BinomiRare).
3. Conduct stratified analysis in parallel with combined analysis. This may be important to detect variants associated with the outcome in only one stratum (e.g. only in females or only in African Americans), or that have larger effect size in one stratum. Aggregated data testing may reduce power to detect such as association.

Finally, we make the additional notes, that do not lead to specific rules but should be considered when planning analyses:

a. When combining diverse studies together, the type 1 error control of the Score test depends on the arrangement of case proportion and carrier numbers across the studies. However, for SPA and BinomiRare there is no such pattern, and only the number of carriers in the combined sample is important.
b. For a fixed number of carriers, power is highly affected by the proportion of cases. If you are interested in limiting the number of tested variants to those with higher statistical power, consider restricting the analysis to variants with high number of carriers in the study with higher disease proportion.

## Discussion

We performed a simulation study to (primarily) assess the type 1 error control of single-variant association tests for a binary outcome when pooling individual level data from heterogeneous studies. The asymptotic framework of our simulations is such that the number of carriers of a variant and the sample sizes were fixed. Variability came from the number of diseased individuals, aka, cases in the sample in general and in the carriers specifically. Despite the fact that testing of rare variant associations suffers from low power, it is still routinely performed as part of genome-wide association studies applied on sequencing or imputed genotype data, or when fine-mapping genomic regions, and investigators need to know when such tests are statistically valid. Our simulations were performed in simplified settings combining two studies, in the absence of covariates, other than study-specific intercept. This allowed for computational efficiency when running a very large number of simulations, and for reducing the potential number of scenarios to investigate. We found that performance of the Score test largely depends on the number of carriers of the rare variant, number of diseased carriers, and the proportion of diseased individuals in the sample. This is, by design, true for the BinomiRare test as well. Notably, the performance of the tests also depends on the properties of the two combined studies: when the combined sample has a fixed number of carriers, test performance differs according to the number of carriers in each of the combined studies, and the outcome prevalence in each.

As was shown in the past, the Score test controls the type 1 error when the case-control proportion is 1. Like other tests, it is conservative when the number of carriers is very low. In simulations combining two studies of equal sizes, with a total number of 110 carriers, even when there were 50% cases in one of the studies and 100 carriers in that study, type 1 error was controlled regardless of the disease prevalence in the study with 10 carriers (Figure 1, row “Score Test”, third column from the left). In these simulations, it seemed like using SPA to re-compute p-values produced better calibration (Figure 1 row “SPA Test”, third column from the left). However, in the simulations with very low number of carriers (10 to 30 in each of the studies), when the two studies had 50% cases, the SPA test often did not control the type 1 error, while the usual Score test did (Supplementary Figure 4). When the case proportion is lower, it is clear that as the number of carriers grows the Score test’s control of type 1 error improves. However, we could not point to a single and simple rule, for when it is appropriate to use the Score test.

Given that the Score test controls the type 1 error when there are 50% cases in each of the studies, a natural question is whether it is useful to sample controls to generate a dataset with 50% cases. As we saw in simulations, this indeed led to control of type 1 error, however also to a loss of power. The idea behind down-sampling of controls is that most of the information is in the cases, and therefore, the loss of information is low. However, as our simulations show, this is not correct. Down-sampling of controls leads to substantial reduction in the sample size, and therefore to both reduction in the quality of estimation of disease probability model, and to reduction in the number of variant carriers used. The properties of the tests depend on the number of carriers.

BinomiRare could be applied with either the usual p-value, or the mid-p-value. While in our primary simulations where the mid-p-value mostly controlled the type 1 error, and almost always improved upon the SPA when it was inflated, we found that the mid-p-value did not control the type 1 error in some settings, especially when the cases were 50% of the sample in both studies, similarly to the SPA. The usual p-value always controlled the type 1 error. Mid-p-values are preferred (when controlling the type 1 error) because they are less conservative. In all, we recommend using the mid-p-values when the case proportion is lower than 50%, and to compute and report both types of p-values.

We formulated a set of recommendations for investigators in studies such as TOPMed, combining together individual-level data from multiple heterogeneous studies. The recommendations take into account the availability of tests across software packages, type 1 error control across extreme settings combining two heterogeneous studies, and power based on modelling assumption and a small number of simulations in the rare variant settings. We did not assess (a) all possible combinations of two studies in terms of their disease proportion and carrier counts, (b) more than two studies, (c) additive mode of inheritance for slightly higher count variants, (d) p-value thresholds lower than 1 × 10^−4^ across all settings, (e) estimation of power while accounting for type 1 error of each test (e.g. by identifying and using the specific p-value threshold yielding the desired type 1 error rate in the power simulations). While doing all these would have been helpful, this is not feasible. Both the number of simulations and the disc memory required for saving a lot of data in order to perform additional computations, would be prohibitive. Therefore, we base our recommendations on simplified and extreme settings of type 1 error control. For power, we primarily show that for rare variants, where dominant mode of inheritance is appropriate as the vast majority of individuals are heterozygotes, the BinomiRare test, which is valid, has often similar performance to the SPA test when it is valid as well, or that SPA test has slightly higher power. For higher frequency variants having homozygotes as well, standard statistical thinking posits that the Score and SPA test will be even more powerful (when valid) because they use additional information, and this is seen to some extent in our simulations as well. Further, while theoretically the BinomiRare test is computationally efficient (in terms of both computer time and memory), current implementations of the Score test use various approaches to speed up matrix computations, making it very efficient in practice, so that BinomiRare is slower. Therefore, when it is valid, the score test is most desirable. The SPA test is less efficient, and its p-value computation is implemented by software packages as re-computation of the naïve Score p-value when it is <0.05. Similar approach could be taken if using BinomiRare test. Still, SPA test is preferred over BinomiRare when it controls type 1 error, due to better power. An important conclusion is that when using the SPA test, our simulations suggest that we can control the total number of carriers rather than the respective number of carriers in each study, i.e. by requiring at least 90-110 carriers in the combined sample. It is not clear what the appropriate minimum number of carriers is (see Figure 3), and it changes by study characteristics. Therefore, it is critically important to perform replication or other follow-up analysis, as is common in genetic association studies. While most of our simulations were performed at p-value threshold of 10^−4^, we also considered lower thresholds for some of the simulations. The conclusions remain the same.

While this work is focused on performance of statistical tests when pooling together data from heterogeneous studies, it highlights issues that are worth addressing in future work. The level of inflation/deflation of the tests when applied on variants with very low number of carriers, varies in different settings (see for example Supplementary Figure 4). Therefore, to assess overall patterns of inflation/deflation due to population stratification, one may need to rely on results from testing common variants, in which tests follow their asymptotic properties. To generate QQ-plots comparing the observed versus the expected distributions of test results when testing rare variants, Lee et al. (2016), developed a resampling based procedure. It would be useful to extend their approach to other tests.

## Author contributions

TS conceptualized and designed the study, performed simulation studies and drafted the manuscript. NG developed data visualization. Both authors critically reviewed the approved the manuscript.

## Acknowledgements

The authors thank Seunggeun Lee, Rounak Dey, and the anonymous reviewers for reviewing the manuscript draft and providing helpful comments. T.S. was supported by grants from National Heart, Lung, and Blood Institute (NHLBI; 1R35HL135818, and 1R21HL145425).

## Data Availability Statement

not applicable.

## Figures

**Supplementary Figure 1:**
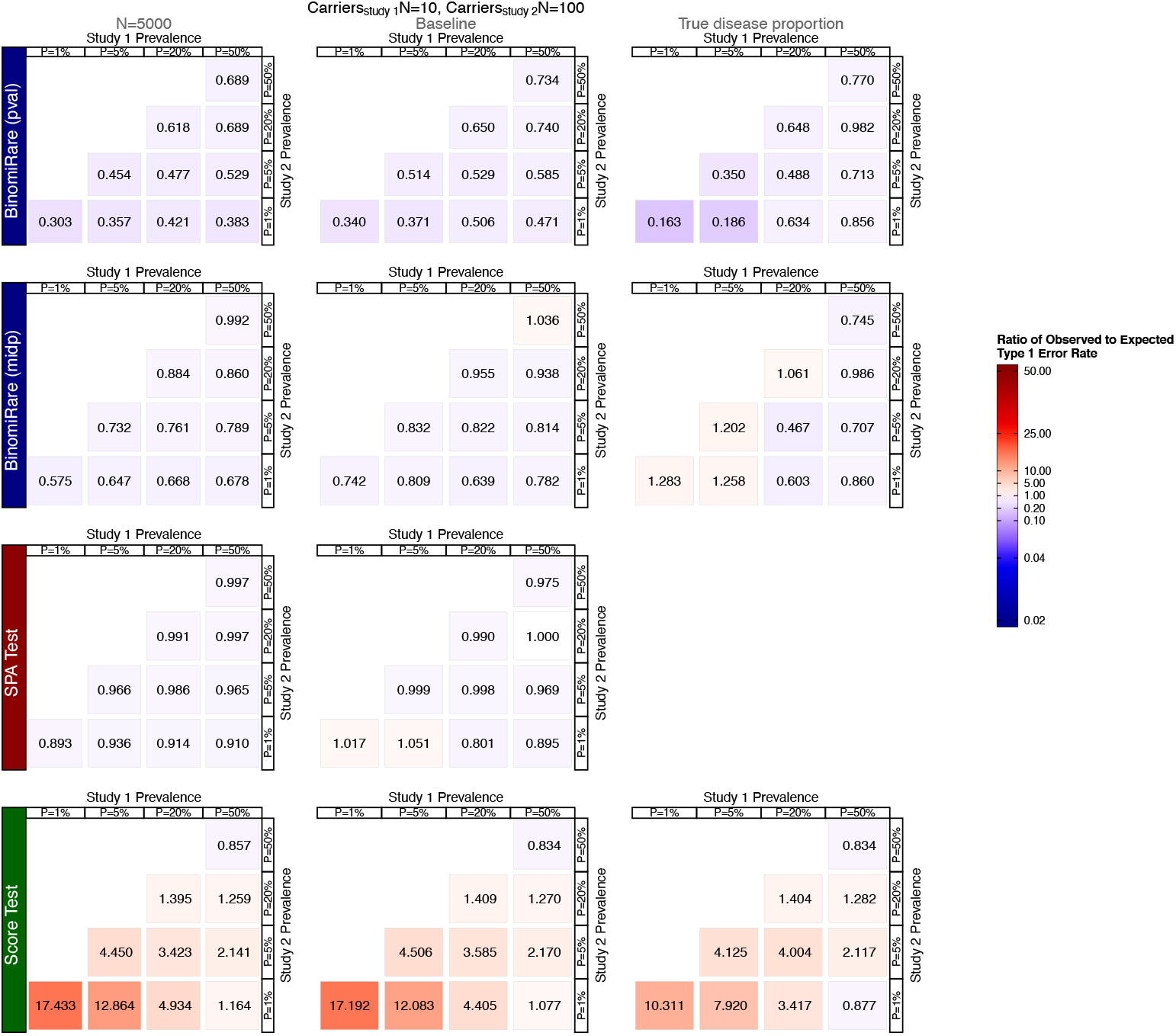
Ratios between observed and expected type 1 error rate in simulation studies when testing a binary outcome for association with a rare genetic variant. We compared the naïve score (Score) test, the SPA test, and BinomiRare with the usual p-values (pval) and the mid-p-value (midp). In all settings data from two studies were pooled together, with 10 carriers of the rare variant in study 1, and 100 carriers of the rare variant in study 2. The left column (“N=5000”) corresponds to settings with 5,000 individuals in each of the studies. The middle column (“Baseline”) corresponds to settings with 10,000 individuals in each of the studies, and the right column (“True disease proportion”) provides results for analyses that plugged-in the true outcome prevalence in each of the studies in the Score statistic (for the Score test) and provided them the BinomiRare test. For each setting we performed 10^8^ simulations, and the p-value threshold used for determining significance was 10^−4^. Values of 1 correspond to perfect calibration, and values larger (smaller) than 1 correspond to inflation (deflation), or higher (lower) number of detected false associations.

**Supplementary Figure 2:**
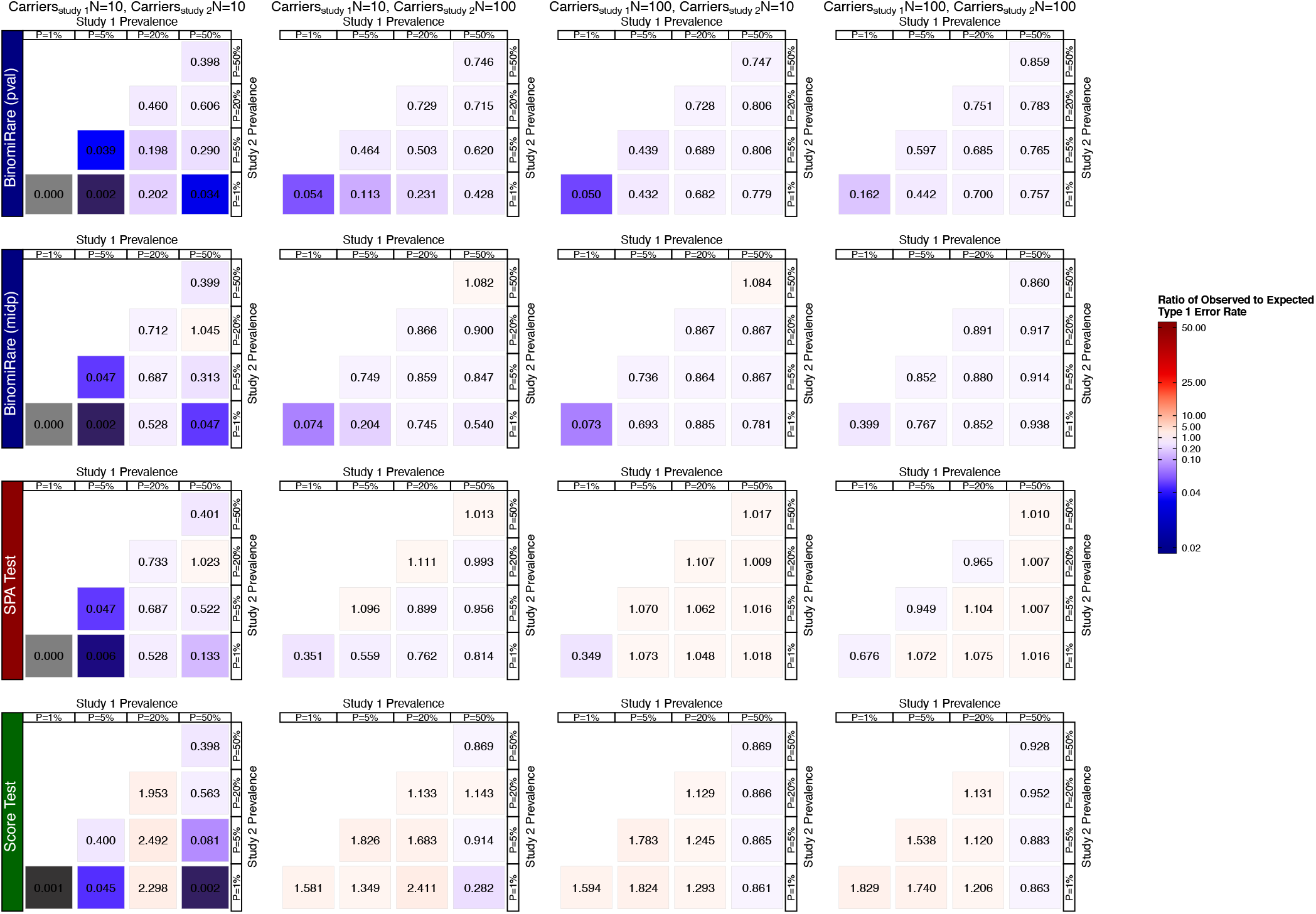
Ratios between observed and expected type 1 error rate in simulation studies when testing a binary outcome for association with a rare genetic variant, and when reducing the sample size by sampling controls to generate samples with up to three controls per case. We compared the naïve score (Score) test, the SPA test, and BinomiRare with the usual p-values (pval) and the mid-p-value (midp), for settings defined by the number of carriers and outcome prevalence in each study. Both studies had n=10,000 individuals before sampling of controls. Controls were sampled in each study separately. For each setting we performed 10^8^ simulations, and the p-value threshold used for determining significance was 10^−4^. Values of 1 correspond to perfect calibration, and values larger (smaller) than 1 correspond to inflation (deflation), or higher (lower) number of detected false associations.

**Supplementary Figure 3:**
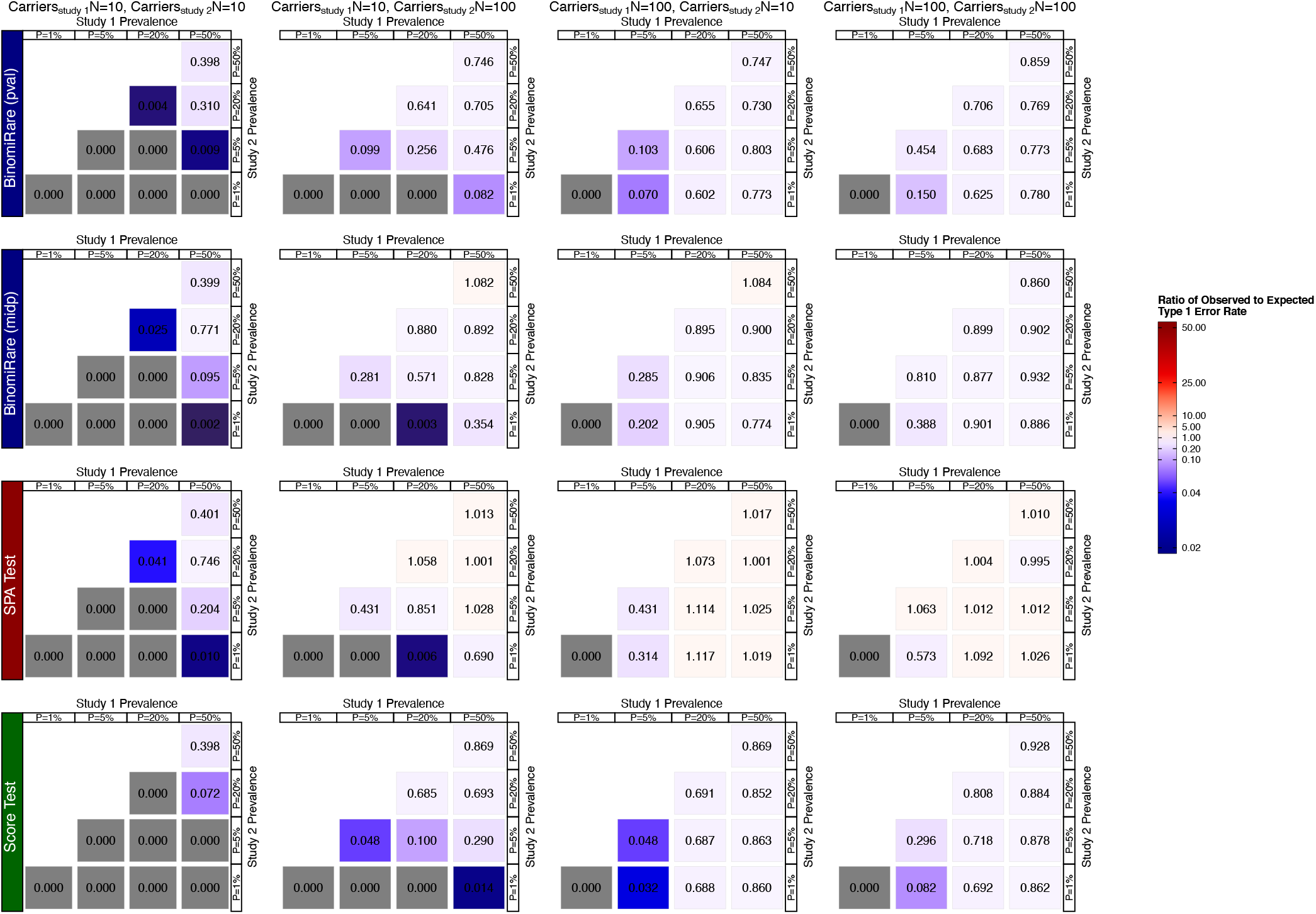
Ratios between observed and expected type 1 error rate in simulation studies when testing a binary outcome for association with a rare genetic variant, and when reducing the sample size by sampling controls to generate samples with one control per case. We compared the naïve score (Score) test, the SPA test, and BinomiRare with the usual p-values (pval) and the mid-p-value (midp), for settings defined by the number of carriers and outcome prevalence in each study. Both studies had n=10,000 individuals before sampling of controls. Controls were sampled in each study separately. For each setting we performed 10^8^ simulations, and the p-value threshold used for determining significance was 10^−4^. Values of 1 correspond to perfect calibration, and values larger (smaller) than 1 correspond to inflation (deflation), or higher (lower) number of detected false associations.

**Supplementary Figure 4:**
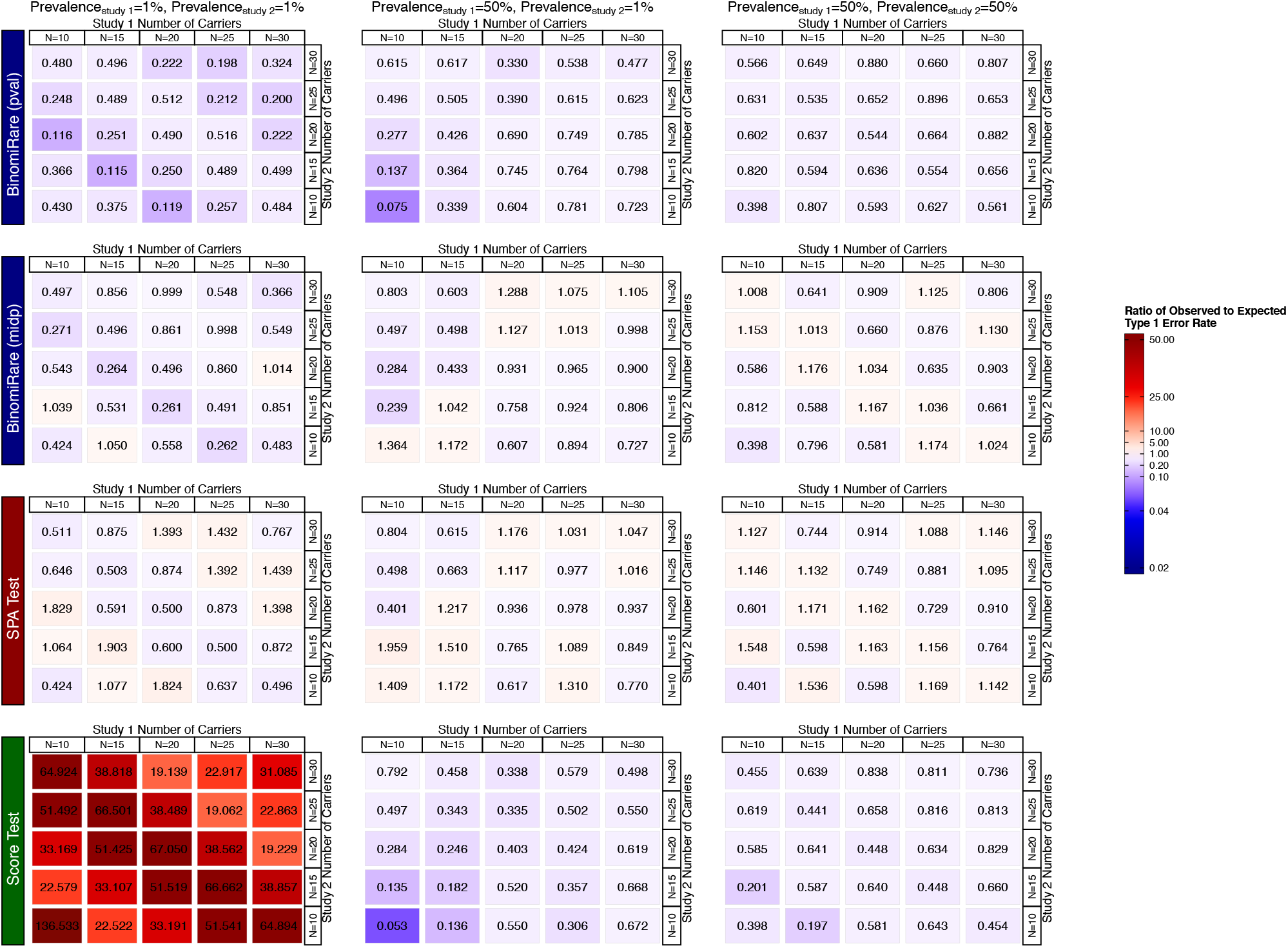
Ratios between observed and expected type 1 error rate in simulation studies when testing a binary outcome for association with a rare genetic variant. We compared the naïve score (Score) test, the SPA test, and BinomiRare with the usual p-values (pval) and the mid-p-value (midp), for settings defined by the number of carriers and outcome prevalence in each study. Both studies had n=10,000 individuals. For each setting we performed 10^8^ simulations, and the p-value threshold used for determining significance was 10^−4^. Values of 1 correspond to perfect calibration, and values larger (smaller) than 1 correspond to inflation (deflation), or higher (lower) number of detected false associations.

**Supplementary Figure 5:**
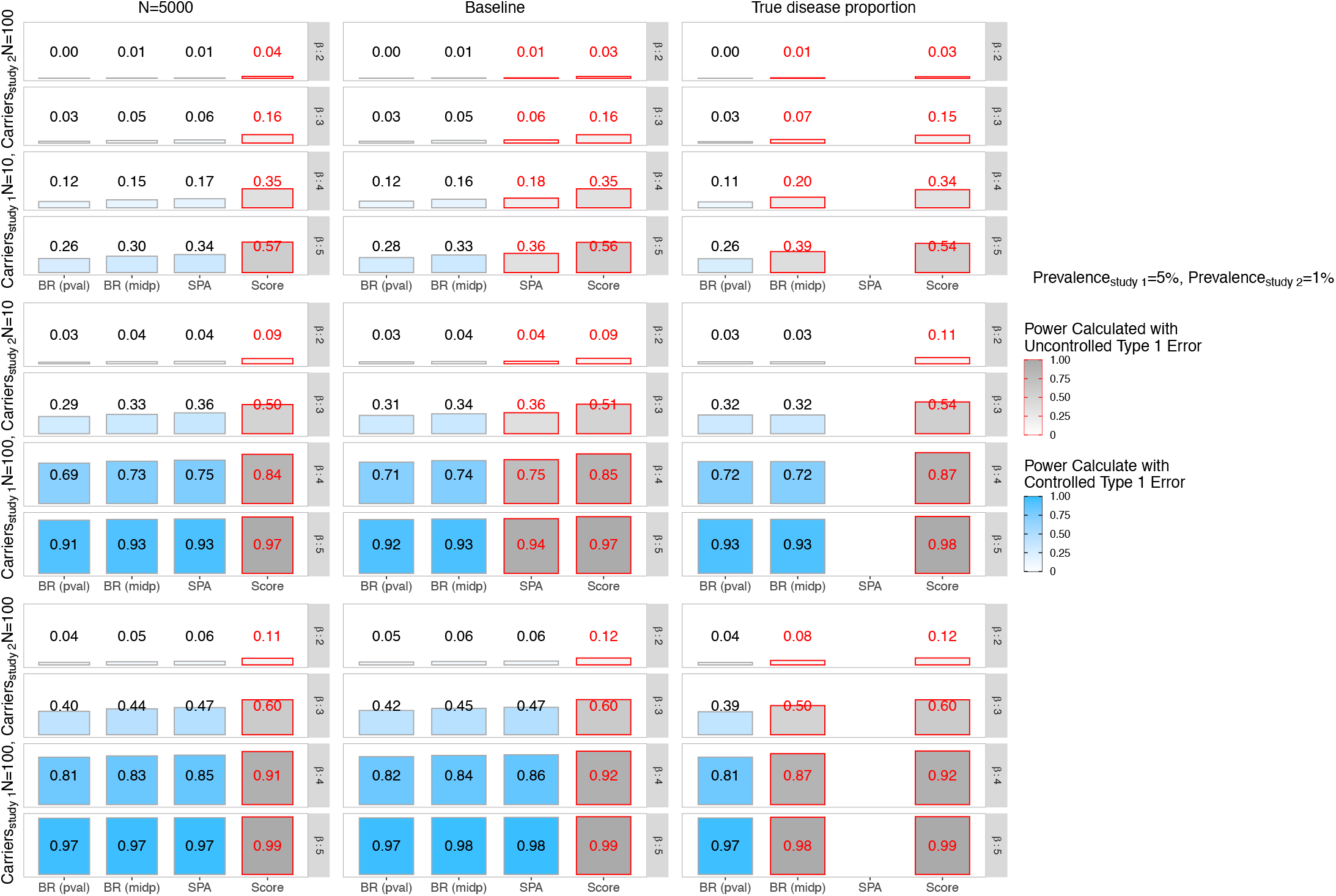
Power estimated in simulation studies when testing a binary outcome for association with a rare genetic variant. We compared the naïve score (Score) test, the SPA test, and BinomiRare with the usual p-values (pval) and the mid-p-value (midp). The simulation settings are defined by the number of carriers in each of the studies, and the variant effect size *β*. The outcome prevalence was fixed at 0.05 in study 1 and 0.01 in study 2. The left column (“N=5000”) corresponds to simulations with 5,000 observations in each study. The middle (“Baseline”) corresponds to simulations with 10,000 observations in each study. The right column (“True disease proportion”) provides results for analyses that plugged-in the true outcome prevalence in each of the studies in the Score statistic (for the Score test) and provided them the BinomiRare test. For each setting we performed 10^4^ simulations, and the p-value threshold used for determining significance was 10^−4^. We color coded the settings according to type 1 error control in the simulations corresponding to the same prevalence, and carrier settings, but with no variant association.

**Supplementary Figure 6:**
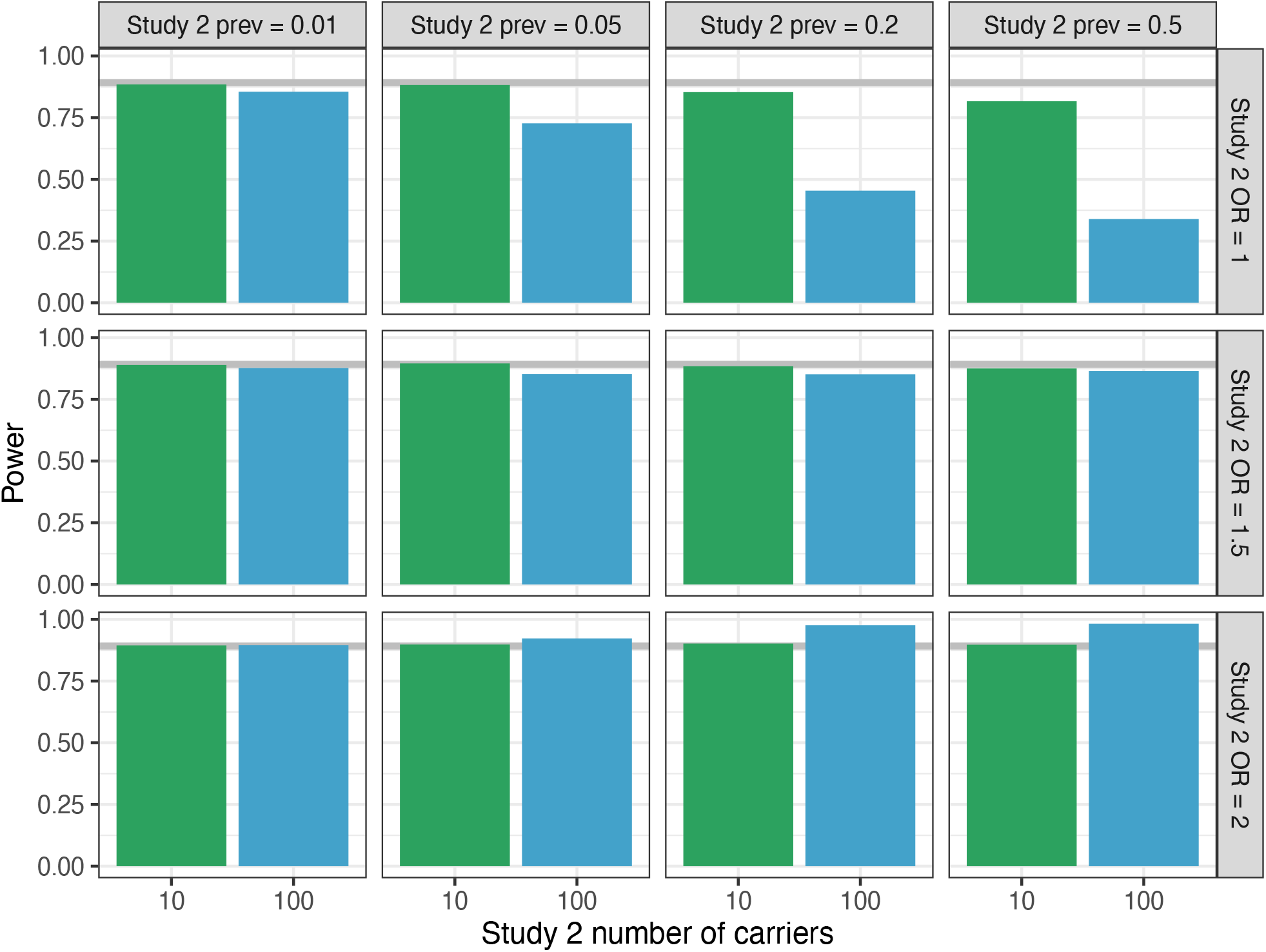
Power estimated in simulation studies when testing a binary outcome for an association with a rare genetic variant using the BinomiRare test (using the usual p-values, pval). In study 1, the outcome prevalence was always 0.5, there were 100 carriers, and the variant effect size was *β*_1*g*_ = log (4). Each block in the figure provides estimated testing power for a given outcome prevalence in study 2 (columns), odds ratio (OR) exp (*β*_2*g*_) in study 2 (rows), and for 10 and 100 carriers in study 2. The grey line corresponds to the estimated power for testing the variant association in study 1 alone. For each setting we performed 10^4^ simulations, and the p-value threshold used for determining significance was 10^−4^.

## Supplementary Tables

**Supplementary Table 1:**
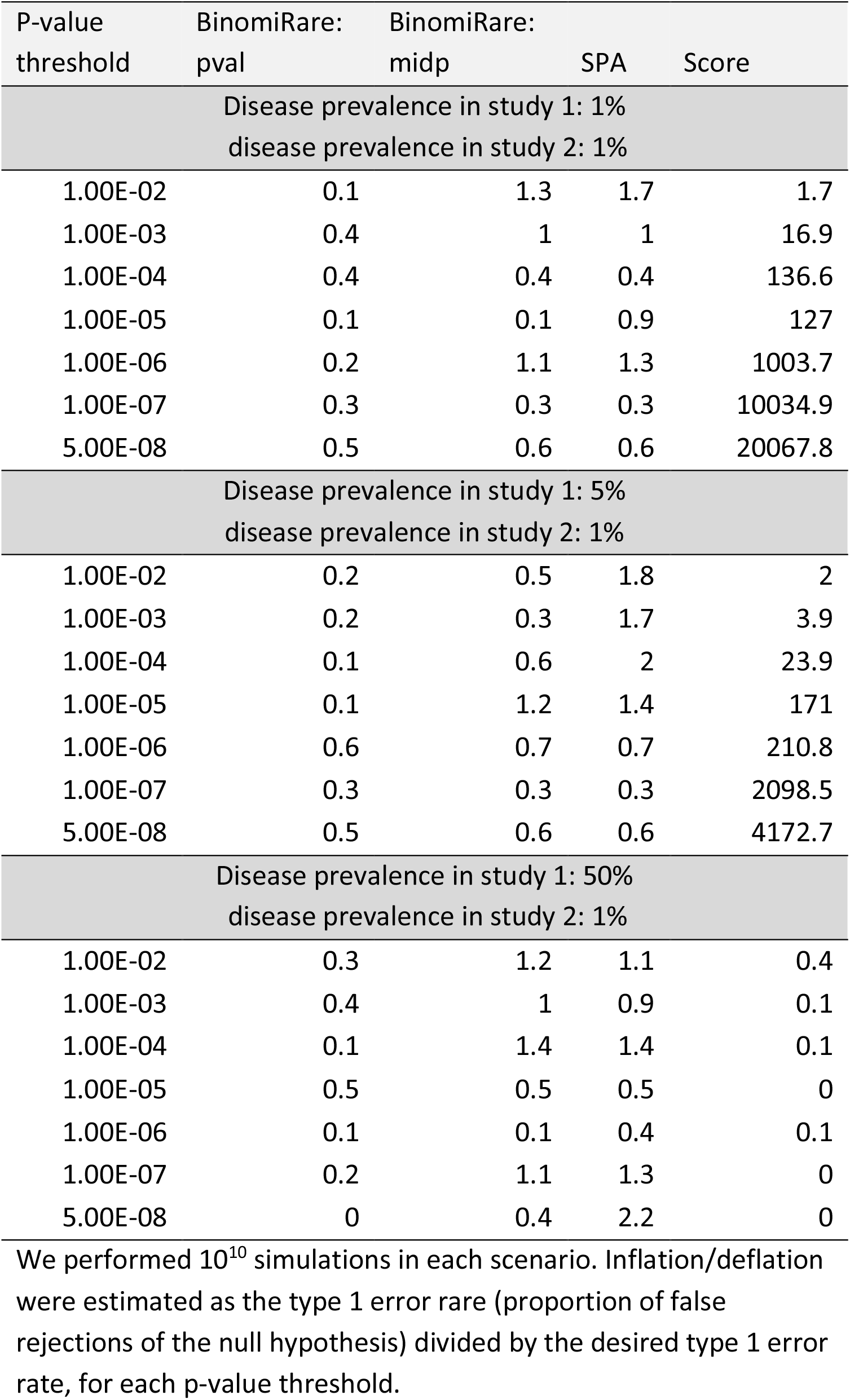
Estimated inflation/deflation of tests of rare variant association with a binary trait, with 10 carriers in each of study 1 and study 2.

| P-value threshold | BinomiRare: pval | BinomiRare: midp | SPA | Score |
| --- | --- | --- | --- | --- |
| Disease prevalence in study 1: 1%<br>disease prevalence in study 2: 1% |  |  |  |  |
| 1.00E-02 | 0.1 | 1.3 | 1.7 | 1.7 |
| 1.00E-03 | 0.4 | 1 | 1 | 16.9 |
| 1.00E-04 | 0.4 | 0.4 | 0.4 | 136.6 |
| 1.00E-05 | 0.1 | 0.1 | 0.9 | 127 |
| 1.00E-06 | 0.2 | 1.1 | 1.3 | 1003.7 |
| 1.00E-07 | 0.3 | 0.3 | 0.3 | 10034.9 |
| 5.00E-08 | 0.5 | 0.6 | 0.6 | 20067.8 |
| Disease prevalence in study 1: 5%<br>disease prevalence in study 2: 1% |  |  |  |  |
| 1.00E-02 | 0.2 | 0.5 | 1.8 | 2 |
| 1.00E-03 | 0.2 | 0.3 | 1.7 | 3.9 |
| 1.00E-04 | 0.1 | 0.6 | 2 | 23.9 |
| 1.00E-05 | 0.1 | 1.2 | 1.4 | 171 |
| 1.00E-06 | 0.6 | 0.7 | 0.7 | 210.8 |
| 1.00E-07 | 0.3 | 0.3 | 0.3 | 2098.5 |
| 5.00E-08 | 0.5 | 0.6 | 0.6 | 4172.7 |
| Disease prevalence in study 1: 50%<br>disease prevalence in study 2: 1% |  |  |  |  |
| 1.00E-02 | 0.3 | 1.2 | 1.1 | 0.4 |
| 1.00E-03 | 0.4 | 1 | 0.9 | 0.1 |
| 1.00E-04 | 0.1 | 1.4 | 1.4 | 0.1 |
| 1.00E-05 | 0.5 | 0.5 | 0.5 | 0 |
| 1.00E-06 | 0.1 | 0.1 | 0.4 | 0.1 |
| 1.00E-07 | 0.2 | 1.1 | 1.3 | 0 |
| 5.00E-08 | 0 | 0.4 | 2.2 | 0 |

We performed $10^{10}$ simulations in each scenario. Inflation/deflation were estimated as the type 1 error rate (proportion of false rejections of the null hypothesis) divided by the desired type 1 error rate, for each p-value threshold.

**Supplementary Table 2:** Estimated inflation/deflation of tests of rare variant association with a binary trait, with 10 carriers in study 1 and 100 carriers in study 2. *Here disease prevalence is always equal to or higher in the study with less carriers* compared to the one with less.

| P-value threshold | BinomiRare: pval | BinomiRare: midp | SPA | Score |
| --- | --- | --- | --- | --- |
| Disease prevalence in study 1: 1%<br>disease prevalence in study 2: 1% |  |  |  |  |
| 1.00E-02 | 0.5 | 0.6 | 0.9 | 2.1 |
| 1.00E-03 | 0.5 | 0.8 | 0.9 | 5.1 |
| 1.00E-04 | 0.3 | 0.8 | 1 | 17.2 |
| 1.00E-05 | 0.3 | 0.6 | 1 | 69.4 |
| 1.00E-06 | 0.3 | 0.6 | 1 | 247.1 |
| 1.00E-07 | 0.2 | 0.5 | 0.9 | 1048.2 |
| 5.00E-08 | 0.3 | 0.4 | 0.8 | 1660.3 |
| Disease prevalence in study 1: 5%<br>disease prevalence in study 2: 1% |  |  |  |  |
| 1.00E-02 | 0.4 | 0.8 | 1.2 | 1.7 |
| 1.00E-03 | 0.5 | 0.7 | 0.8 | 4 |
| 1.00E-04 | 0.4 | 0.8 | 1.1 | 12.1 |
| 1.00E-05 | 0.3 | 0.7 | 1.1 | 50 |
| 1.00E-06 | 0.3 | 0.6 | 1 | 146.2 |
| 1.00E-07 | 0.3 | 0.5 | 0.9 | 635.1 |
| 5.00E-08 | 0.3 | 0.5 | 0.9 | 899.2 |
| Disease prevalence in study 1: 50%<br>disease prevalence in study 2: 1% |  |  |  |  |
| 1.00E-02 | 0.7 | 0.9 | 1.3 | 1.3 |
| 1.00E-03 | 0.7 | 0.8 | 0.9 | 0.8 |
| 1.00E-04 | 0.5 | 0.8 | 0.9 | 1.1 |
| 1.00E-05 | 0.4 | 0.8 | 1 | 1.7 |
| 1.00E-06 | 0.3 | 0.6 | 1 | 3.3 |
| 1.00E-07 | 0.3 | 0.6 | 1 | 7.5 |
| 5.00E-08 | 0.3 | 0.5 | 0.9 | 6.7 |

We performed $10^{10}$ simulations in each scenario. Inflation/deflation were estimated as the type 1 error rare (proportion of false rejections of the null hypothesis) divided by the desired type 1 error rate, for each p-value threshold.

**Supplementary Table 3:** Estimated inflation/deflation of tests of rare variant association with a binary trait, with 100 carriers in study 1 and 10 carriers in study 2. *Here disease prevalence is always equal to or higher in the study with more carriers* compared to the one with less.

| P-value threshold | BinomiRare: pval | BinomiRare: midp | SPA | Score |
| --- | --- | --- | --- | --- |
| Disease prevalence in study 1: 1%<br>disease prevalence in study 2: 1% |  |  |  |  |
| 1.00E-02 | 0.5 | 0.6 | 0.9 | 2.1 |
| 1.00E-03 | 0.5 | 0.8 | 0.9 | 5.1 |
| 1.00E-04 | 0.3 | 0.8 | 1 | 17.2 |
| 1.00E-05 | 0.3 | 0.6 | 1 | 69.3 |
| 1.00E-06 | 0.3 | 0.6 | 1 | 247 |
| 1.00E-07 | 0.2 | 0.5 | 0.9 | 1047.7 |
| 5.00E-08 | 0.3 | 0.5 | 0.9 | 1659.8 |
| Disease prevalence in study 1: 5%<br>disease prevalence in study 2: 1% |  |  |  |  |
| 1.00E-02 | 0.6 | 1 | 1 | 1.1 |
| 1.00E-03 | 0.5 | 0.8 | 1 | 2.1 |
| 1.00E-04 | 0.5 | 0.8 | 1 | 4.9 |
| 1.00E-05 | 0.5 | 0.8 | 1 | 11.9 |
| 1.00E-06 | 0.4 | 0.7 | 1 | 30.4 |
| 1.00E-07 | 0.4 | 0.6 | 0.9 | 81.7 |
| 5.00E-08 | 0.4 | 0.6 | 0.9 | 110.3 |
| Disease prevalence in study 1: 50%<br>disease prevalence in study 2: 1% |  |  |  |  |
| 1.00E-02 | 0.9 | 0.9 | 1 | 1 |
| 1.00E-03 | 0.9 | 0.9 | 1 | 0.9 |
| 1.00E-04 | 0.8 | 0.8 | 1 | 0.8 |
| 1.00E-05 | 0.8 | 0.8 | 1 | 0.7 |
| 1.00E-06 | 0.6 | 1 | 1 | 0.6 |
| 1.00E-07 | 0.6 | 1 | 1 | 0.5 |
| 5.00E-08 | 0.6 | 1.1 | 1 | 0.5 |

We performed $10^{10}$ simulations in each scenario. Inflation/deflation were estimated as the type 1 error rate (proportion of false rejections of the null hypothesis) divided by the desired type 1 error rate, for each p-value threshold.

**Supplementary Table 4:**
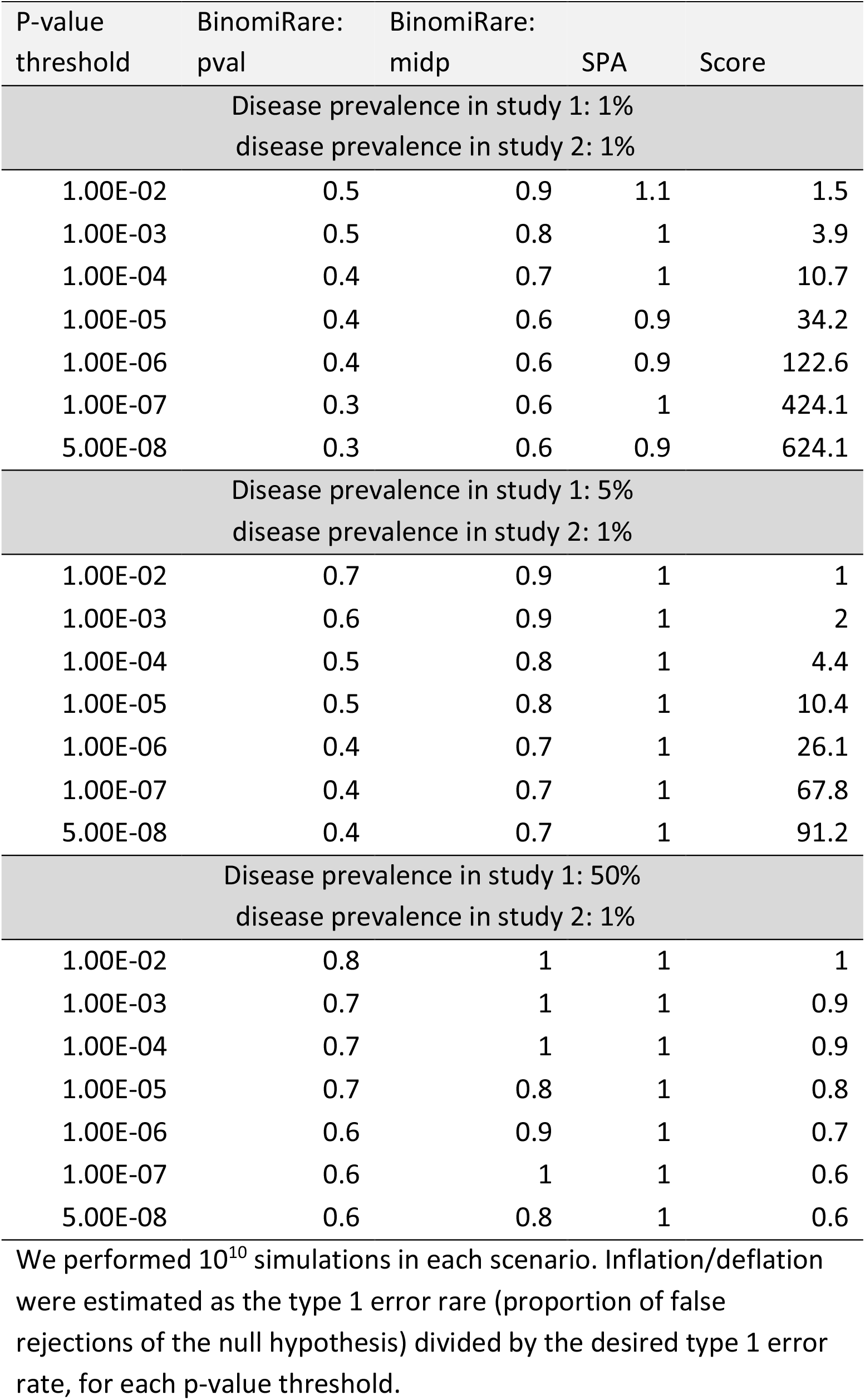
Estimated inflation/deflation of tests of rare variant association with a binary trait, with 100 carriers in both studies.

| P-value threshold | BinomiRare: pval | BinomiRare: midp | SPA | Score |
| --- | --- | --- | --- | --- |
| Disease prevalence in study 1: 1%<br>disease prevalence in study 2: 1% |  |  |  |  |
| 1.00E-02 | 0.5 | 0.9 | 1.1 | 1.5 |
| 1.00E-03 | 0.5 | 0.8 | 1 | 3.9 |
| 1.00E-04 | 0.4 | 0.7 | 1 | 10.7 |
| 1.00E-05 | 0.4 | 0.6 | 0.9 | 34.2 |
| 1.00E-06 | 0.4 | 0.6 | 0.9 | 122.6 |
| 1.00E-07 | 0.3 | 0.6 | 1 | 424.1 |
| 5.00E-08 | 0.3 | 0.6 | 0.9 | 624.1 |
| Disease prevalence in study 1: 5%<br>disease prevalence in study 2: 1% |  |  |  |  |
| 1.00E-02 | 0.7 | 0.9 | 1 | 1 |
| 1.00E-03 | 0.6 | 0.9 | 1 | 2 |
| 1.00E-04 | 0.5 | 0.8 | 1 | 4.4 |
| 1.00E-05 | 0.5 | 0.8 | 1 | 10.4 |
| 1.00E-06 | 0.4 | 0.7 | 1 | 26.1 |
| 1.00E-07 | 0.4 | 0.7 | 1 | 67.8 |
| 5.00E-08 | 0.4 | 0.7 | 1 | 91.2 |
| Disease prevalence in study 1: 50%<br>disease prevalence in study 2: 1% |  |  |  |  |
| 1.00E-02 | 0.8 | 1 | 1 | 1 |
| 1.00E-03 | 0.7 | 1 | 1 | 0.9 |
| 1.00E-04 | 0.7 | 1 | 1 | 0.9 |
| 1.00E-05 | 0.7 | 0.8 | 1 | 0.8 |
| 1.00E-06 | 0.6 | 0.9 | 1 | 0.7 |
| 1.00E-07 | 0.6 | 1 | 1 | 0.6 |
| 5.00E-08 | 0.6 | 0.8 | 1 | 0.6 |

We performed $10^{10}$ simulations in each scenario. Inflation/deflation were estimated as the type 1 error rare (proportion of false rejections of the null hypothesis) divided by the desired type 1 error rate, for each p-value threshold.

## Notes

### Competing Interest Statement

The authors have declared no competing interest.

### Summary of Updates

We have added simulations looking at power when pooling together two studies in which the variant effect size differ between the studies; added simulations looking at lower p-value thresholds; updated the recommendations.

